# A neonicotinoid pesticide causes tissue-specific gene expression changes in bumble bees

**DOI:** 10.1101/2024.10.09.612896

**Authors:** Alicja Witwicka, Federico López-Osorio, Hannah Chaudhry-Phipps, Yannick Wurm

## Abstract

Pesticides often harm beneficial insect pollinators, impairing their ability to navigate the environment, learn, fight off disease, and reproduce. Understanding the mechanisms behind these disorders is essential for improving pesticide risk assessments. To test whether pesticide exposure induces similar or distinct transcriptional responses across tissues, we administered field-realistic dose of the common neonicotinoid clothianidin to *Bombus terrestris* bumble bees. We then measured gene expression in brains, hind femurs, and Malpighian tubules. Our analyses revealed that 82% of gene expression differences were tissue-specific. Although genes associated with energy metabolism were consistently down-regulated across all tissues, pesticide exposure primarily affected core tissue functions, namely genes linked to ion transport in the brain, muscle function in the hind femur, and detoxification in Malpighian tubules. Furthermore, while the brain holds the highest abundance of pesticide target receptors, other tissues showed more substantial differences in gene expression magnitude. These findings reveal that pesticide exposure causes complex, tissue-specific effects rather than a uniform body-wide response. Our study provides a mechanistic basis for the severe effects of pesticide exposure on bees and shows how transcriptomics can help pinpoint the most affected areas and processes across the body. Accordingly focusing toxicological assays could significantly improve the precision of pesticide safety evaluations.

## Introduction

Ensuring the health of insect pollinators is essential for ecosystem stability and global food security (Potts et al., 2016; Wagner et al., 2021). Although various hazards threaten these insects, pesticide exposure poses a notable risk in agricultural settings worldwide (Goulson et al., 2015; Raine & Rundlöf, 2024). Indeed, numerous studies have demonstrated pesticides cause unintended harm to pollinating insects (Wood & Goulson, 2017). Thus, understanding how broad-spectrum pesticides impair these insects is crucial for improving pesticide safety assessments.

Our understanding of how pesticides cause damage remains limited despite growing evidence of risks. Previous studies on insect pollinators have demonstrated that pesticides impair brain growth during development (Smith et al., 2020), decrease mandibular gland size (Kozii et al., 2021), reduce sperm mobility (Chaimanee et al., 2016; Strobl et al., 2021), disrupt immunity, and affect macronutrient metabolism (Christen et al., 2018; Colgan et al., 2019; Di Prisco et al., 2013). These findings suggest a general interference of gene regulatory networks across various organs. Identifying and understanding how pesticides disrupt genes and pathways in insect tissues is thus becoming increasingly critical.

Although pesticides propagate throughout organisms, how distinct tissues absorb, break down, and accumulate such substances can vary substantially (Ohtsu et al., 2018). For example, the brain contains an overabundance of the nicotinic acetylcholine receptors targeted by many pesticides, likely rendering the brain more vulnerable (Jones & Sattelle, 2010; Witwicka et al., 2023). Contrarily, foraging honey bees overexpress cytochrome P450s detoxification enzymes in specialized leg structures that carry pollen containing natural plant chemicals and potentially pesticides, likely neutralizing their negative impacts (Mao et al., 2015).

To disentangle the molecular pathways underlying pesticide effects, we analyzed the tissue-specific responses in a common European insect pollinator, the buff-tailed bumble bee *Bombus terrestris*. We hypothesized that pesticide effects would be most severe in the brain and disrupt similar cellular processes across tissues despite their distinct functions. Using transcriptome-wide (RNA-seq) analyses, we measured the consequences of exposure to the neonicotinoid clothianidin in tissues integral for neural processing, motor ability, and waste removal – brain, hind femur, and Malpighian tubules, respectively (Scholer & Krischik, 2014; Smith et al., 2020). We found that clothianidin elicits tissue-specific gene expression profiles connected to core physiological functions. These insights advance our mechanistic understanding of how pesticides harm insect pollinators.

## Results

To investigate the effects of pesticide exposure on the transcriptome across tissues, we analyzed brains, hind femurs, and Malpighian tubules from bumble bees subjected to either control or clothianidin (4.4 ppb) treatments over 12 days (Figure 1A). We used eight replicates for each tissue-treatment combination and examined the expression of 11,874 genes in the bumble bee genome. On average, each replicate generated 34.7 million pairs of 40 bp reads. The dose we administered is representative of pesticide residues found in pollen and nectar (Wood & Goulson, 2017). It does not cause death rapidly but is sufficient to cause phenotypic impacts – clothianidin exposure increased the likelihood of arrested ovarian development by sevenfold compared to the control (standard error = 0.81, Z-value = 2.39, *P* = 0.017, Supplementary Figure 1).

**Figure 1.**
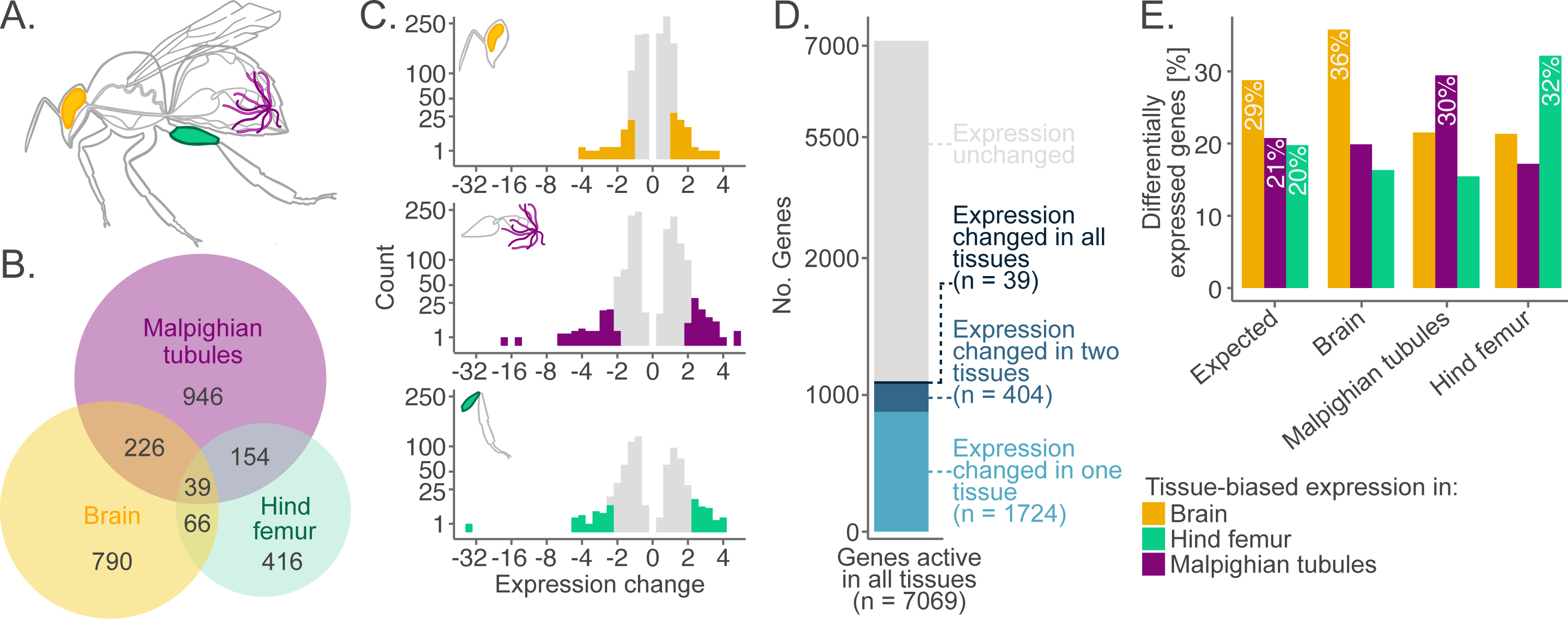
Trends in gene expression changes across tissues. **A.** Schematic representation of tissues used for RNA-seq: brain, hind femur, and Malpighian tubules. **B.** Euler diagram summarizing the distribution and overlap of differentially expressed genes across three tissues. **C.** Histograms showing the distributions of the magnitude of gene expression changes. For each tissue, the 10% of genes with the strongest expression changes are darkened. In Malpighian tubules and hind femur, these genes exhibited at least a two-fold change in expression. However, in the brain, expression changes were below this threshold for most genes. **D.** Classification of genes active in all three tissues. Most of the genes affected by pesticide exposure were differentially expressed in only one tissue despite being active in all tissues. **E.** In each tissue, more of the differentially expressed genes were associated with a cluster of genes with higher (tissue-biased) expression in this particular tissue than would be expected by chance.

### Clothianidin induces tissue-specific changes

Clothianidin exposure induced distinct, tissue-specific transcriptomic changes. Specifically, we found 1365 genes differentially expressed in Malpighian tubules, 1121 in the brain and 675 in the hind femur (Figure 1B). Next, we tested whether these differences between tissues may be because clothianidin-affected genes are only active in some tissues. There was no such effect: most clothianidin-affected genes were still active in all three tissues (Figure 1B and D).

The most highly expressed genes in a tissue contribute substantially to tissue function (Hekselman & Yeger-Lotem, 2020). To test whether such highly expressed genes tend to be more prone to changes in response to clothianidin exposure, we allocated genes into six clusters according to their relative expression levels (898 to 3004 genes per cluster; Supplementary Figure 2). We found that 36% of differentially expressed genes detected in the brain, 32% in the hind femur, and 30% in the Malpighian tubules belonged to clusters exhibiting elevated expression specifically in the respective tissues (Figure 1E). Moreover, these percentages were higher than expected compared to the relative numbers of genes in each cluster (all Z-test *P*-values < 10^-9^). These patterns were even more striking when we considered only the top 10% of genes with the most extreme expression changes, where 50% in the brain, 54% in the hind femur, and 36% in the Malpighian tubules exhibited tissue-specific responses. These results indicate a consistent tissue-specific relationship between high baseline expression and susceptibility to clothianidin exposure. These disruptive effects of pesticide exposure on core tissue functions likely underpin organism-level pathologies and disorders.

### Intensity of clothianidin-induced changes varies between tissues

Intriguingly, the intensity of gene expression changes was uncorrelated with the number of differentially expressed genes. The brain ranked second in the number of differentially expressed genes detected, yet the magnitude of expression changes was less pronounced compared to the other tissues: median gene expression changes in the brain were by 26%, whereas the Malpighian tubules and hind femur exhibited median changes of 50%. Further supporting this pattern, among the top 10% of genes with the strongest expression changes in the brain, most exhibited less than a two-fold change, with the most extreme changes not exceeding four-fold. By contrast, the top 10% of affected genes in the Malpighian tubules and hind femur displayed much more substantial changes, reaching 16-fold and 32-fold, respectively (Figure 1C). This pattern suggests that different tissues may experience pesticide exposure at varying intensities.

### Clothianidin affects pathways critical for tissue function

Pesticide exposure caused distinct changes in biological pathways and processes across tissues. In the hind femur, we observed significant changes in the expression of muscle-specific genes, such as troponin C, myophilin, and muscle-specific protein 20 (Figure 2A). Additionally, lipid transport and localization Gene Ontology terms were enriched in down-regulated genes (Figure 3A), indicating that exposure to clothianidin may compromise muscle function and regeneration. In the Malpighian tubules, Gene Ontology terms linked to DNA helicase activity were enriched, suggesting changes in DNA metabolism and gene regulatory mechanisms, as well as terms associated with ribosomal processes and peptide metabolism (Figure 2B and 3A). The brain exhibited a different profile, including changes in expression of genes that form ion transporters, such as the sodium-independent sulphate anion transporter and the calcium-activated potassium channel slowpoke (Figure 2C). Gene Ontology analysis highlighted the enrichment of genes linked to peptide metabolism, mitochondrial functioning, and transmembrane transport in the brain (Figure 3A).

**Figure 2.**
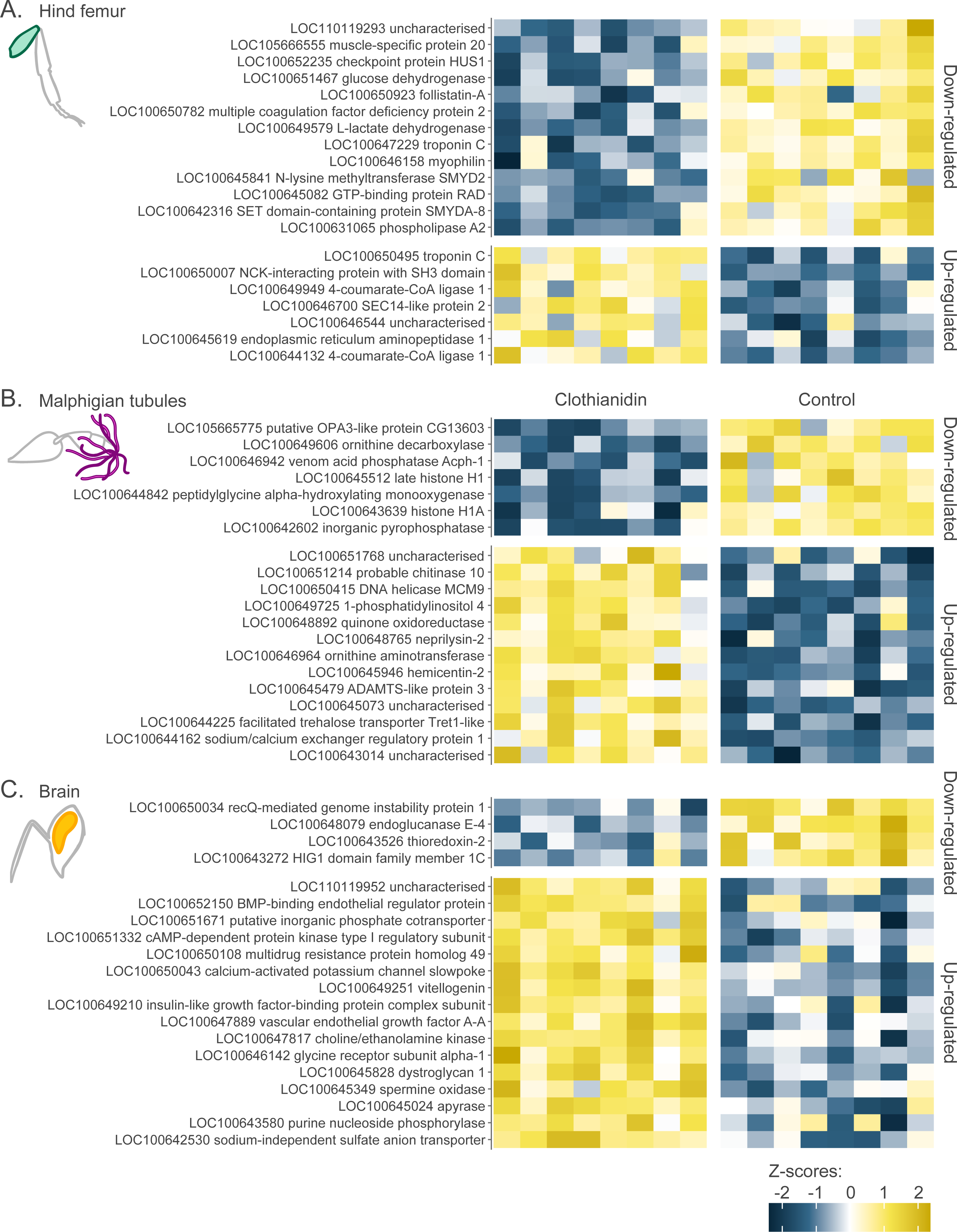
Top 20 differentially expressed genes with the lowest adjusted *P*-values (FDR) for **A.** the hind femur, **B.** Malpighian tubules, and **C.** the brain. All sets of genes were unique for each tissue and contained genes important for their proper functioning. The scale represents z-z-scores scaled across the genes.

**Figure 3.**
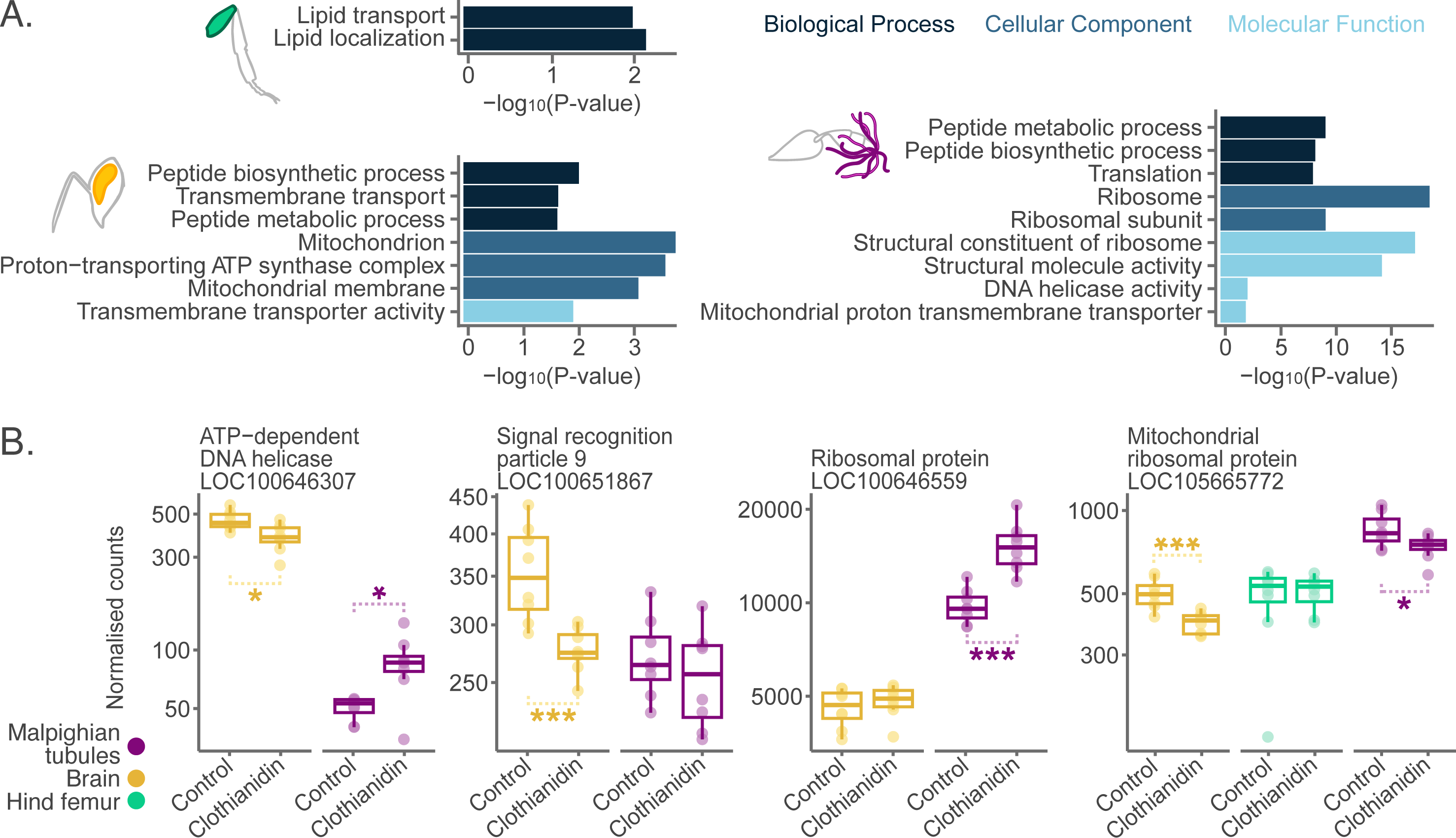
Tissue-specific genes and pathways affected after exposure to clothianidin. **A.** Gene Ontology (GO) terms enriched in differentially expressed genes after clothianidin exposure. Most GO terms were overrepresented in a single tissue. **B.** Boxplots showing selected differentially expressed genes linked to protein metabolism which was enriched in differentially expressed genes mostly up-regulated in the Malpighian tubules and down-regulated int the brain. Significance codes: * < 0.05; ** < 0.01; *** < 0.001.

Tissues across the body also exhibited divergent patterns. For example, ATP−dependent DNA helicase, an enzyme essential in DNA metabolism, was down-regulated in the brain but up-regulated in the Malpighian tubules. The general down-regulation of peptide metabolism in the brain contrasted with its up-regulation in Malpighian tubules, and different genes participating in this process were affected in the two tissues. For example, signal recognition particle 9, important in directing nascent polypeptides from ribosomes to the endoplasmic reticulum, was down-regulated in the brain but unchanged in the Malpighian tubules. While multiple ribosomal proteins were up-regulated in Malpighian tubules, they remained intact in the brain. However, mitochondrial ribosomal proteins were down-regulated in both tissues while remaining unchanged in hind femurs (Figure 3B).

### Clothianidin causes tissue-specific changes in the expression of detoxification genes

The molecular machinery that evolved to counteract naturally occurring xenobiotics also plays a critical role in detoxifying pesticides (Bass et al., 2024). However, the extent to which detoxification processes vary across insect tissues remains underexplored. To detect differences in the expression of detoxification genes between tissues, we identified 120 putative detoxification genes, including 51 cytochrome P450s, 40 ABC-transporters, eleven UDP-glycosyltransferases, twelve glutathione S-transferases, and six detoxification-related carboxylesterases (Supplementary Table 1). Genes from all families were active in each tissue, with cytochrome P450s contributing most to the total expression of detoxification enzymes in the brain and Malpighian tubules. By contrast, the hind femur exhibited similar expression levels of cytochrome P450s and ABC-transporters (Figure 4A).

**Figure 4.**
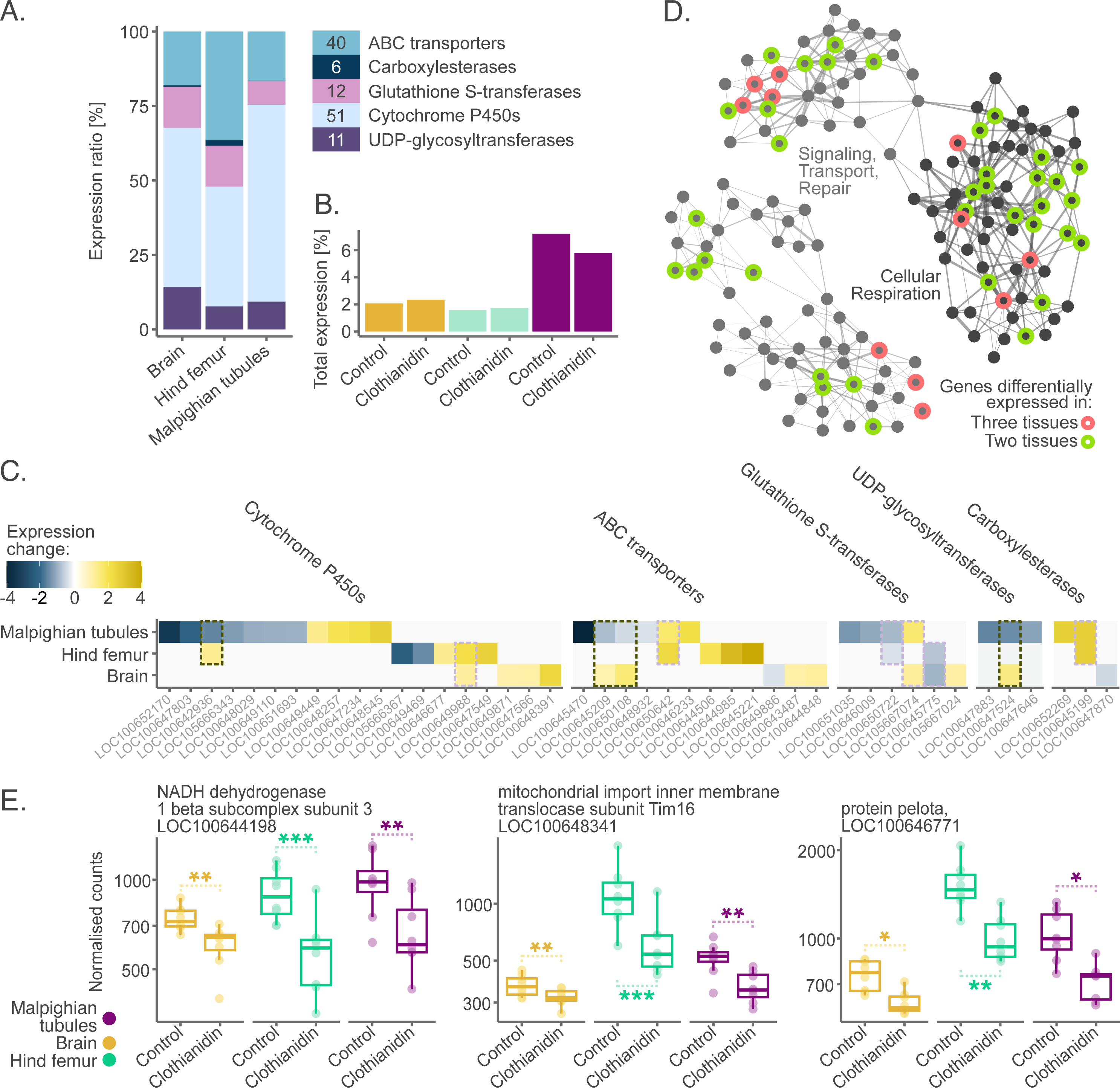
Processes affected across the tissues: detoxification and energy metabolism genes. **A.** Proportions of gene expression from five detoxification gene families in healthy untreated bumble bees. Legend indicates total numbers of genes in each family. **B.** Expression of detoxification genes in the three tissues as a percentage of the total gene expression in the control and clothianidin-exposed samples. After exposure, the overall activity of detoxification genes decreased in the Malpighian tubules and increased in the brain and Malpighian tubules. **C.** Heatmap of expression changes in detoxification genes across the three tissues. There was no systematic direction of expression changes. Furthermore, there was a minimal overlap between tissues with regards to genes affected. Four detoxification genes were up-regulated in one tissue and down-regulated in another (dashed black outlines). **D.** Network analysis of all genes revealed a single module of co-regulated genes affected by clothianidin across all tissues. Each dot represents a gene (n = 153), with line width indicating the correlation level between expression of the genes. Spatial arrangement reflects the number of connections above a specified correlation threshold. We highlight genes significantly differentially expressed in two (light green) and three (pink) tissues. **E.** Examples of differentially expressed genes from the co-regulated module and their expression levels across the control and exposed tissues.

The overall activity of detoxification genes was highest in the Malpighian tubules, consistent with the known role of this organ in detoxification, followed by the brain and hind femur (Figure 4B). Surprisingly, most detoxification genes showing differential expression were unique to a single tissue, with no common detoxification genes differentially expressed across all three tissues (Figure 4C). Intriguingly, while the expression of detoxification genes generally increased in the brain and hind femur, their expression levels decreased overall in the Malpighian tubules (Figure 4B and C). Most changes in detoxification gene expression were moderate, with the majority showing less than two-fold shifts and only the most extreme genes showing four-fold changes (Figure 4C). At least one gene from each detoxification gene family was differentially expressed in all tissues except for UDP-glycosyltransferases, none of which showed differential expression in the hind femur. Some of the differentially expressed genes exhibited relatively high baseline activity in the affected tissue. However, this pattern was not consistent across all genes, indicating that pesticide exposure may not specifically affect highly active detoxification genes (Supplementary Table 1).

### Clothianidin affects energy metabolism in all three tissues

While some genes can be essential across multiple tissues, the local cellular environment, signaling pathways, and physiological needs of individual tissues determine gene activity. To assess the consequences of pesticide exposure on metabolic processes across tissues, we compared the pathways associated with genes differentially expressed in response to clothianidin treatment. Thirty-nine genes were differentially expressed across all three tissues, more than expected by chance (hypergeometric test *P* < 10^-5^, Figure 1B), but only constitutes 1.5% of the total 2,647 differentially expressed genes across all tissues, suggesting limited conserved responses to clothianidin.

To further explore coordinated expression shifts instead of single gene changes, we employed Weighted Gene Co-Expression Network Analysis (WGCNA). This analysis revealed one module comprising 153 genes with correlated gene expression changes across all three tissues following clothianidin exposure (FDR = 0.04, Figure 4C). The eigengene values indicated a general down-regulation of these genes in all tissues after clothianidin exposure. Eleven (28%) of the 39 genes with significant expression changes were part of this module, and all were down-regulated.

The identified module included two functional sub-modules. One sub-module was enriched in genes such as NADH dehydrogenase and cytochrome c oxidase subunit genes (Figure 4D), associated with aerobic and anaerobic respiration, mitochondrial functioning, and ATP production Gene Ontology terms as well as the oxidative phosphorylation KEGG pathway, thus suggesting that clothianidin exposure likely disturbs energy metabolism across the body. No Gene Ontology terms were enriched in the other sub-module. However, this module included genes linked to cellular signaling, transport, and repair, such as mitochondrial transporters or the highly conserved pelota gene, which ensure the integrity and proper functioning of mRNA molecules (Hilal & Spahn, 2015, Figure 4D and E).

## Discussion

Assessing and mitigating the ecological risks of pesticides requires a granular understanding of how these agrochemicals harm beneficial insects. Our study provides insights into tissue-specific transcriptional responses to the neonicotinoid pesticide clothianidin in a widespread bumble bee species. Instead of an even response to pesticide exposure throughout the body, we found highly localized gene expression changes based on tissue-specific physiological roles. Our findings underscore the potential benefits of incorporating differential tissue responses into assessing pesticide risks.

### Baseline tissue metabolism determines the response to pesticide exposure

We demonstrate that low-dose, chronic exposure to the neonicotinoid clothianidin induces tissue-specific transcriptomic changes. Most genes are active in multiple tissues at varying levels, indicating that the same gene can have different roles depending on the tissue (Hekselman & Yeger-Lotem, 2020; Li et al., 2022). Indeed, most genes identified as differentially expressed in only one tissue were still active across all three tissues. However, many of these differentially expressed genes had higher baseline expression levels in the tissues where they were affected. Genes with tissue-biased expression levels likely participate in essential tissue processes and their disruption is thus more likely to cause malfunction or failure. Interestingly, similar tissue-specific alterations of gene expression are present during cancer and aging (dos Santos et al., 2023). The tissue-specific disruption of genes and pathways likely results in more complex exposure outcomes than if a uniform set of processes were affected across the entire insect body.

### Brain tissue may not be the primary target of pesticide effects

Many modern pesticides, including neonicotinoids and sulfoximines specifically target nicotinic acetylcholine receptors (nAChRs), leading to disturbances in synaptic calcium ion (Ca^2+^) levels and neuronal excitability. However, calcium ions also play a critical role in regulating muscle contraction, ATP production, and mitochondrial energy metabolism (Xu et al., 2022). Because baseline metabolism varies across organs, cascading effects of pesticide exposure on a second messenger molecule such as Ca^2+^ may disrupt different metabolic pathways in each tissue.

The brain is often considered a primary target for pesticides due to its overabundance of nAChRs (Jones & Sattelle, 2010; Witwicka et al., 2023). While our data revealed many differentially expressed genes in the brain, the intensity of changes was generally lower than in other tissues. This result challenges the premise that the nervous system is most susceptible to pesticide exposure, adversely affecting learning, memory, and communication (Cabirol & Haase, 2019). In fact, numerous studies have reported learning ability as unaffected by clothianidin in bumble bees (Muth & Leonard, 2019; Piiroinen et al., 2016, 2016; Piiroinen & Goulson, 2016). In insects, glial cells tightly regulate the entry of molecules into the brain, maintaining a stable environment for the neurons, much like the blood-brain barrier in mammals (Limmer et al., 2014). Moreover, neurons can restore their baseline function despite ongoing exposure to stressors through readjustments of ion channel expression and neurotransmitter release (Davis & Müller, 2015). Possibly, the disruption of genes that encode ion transporters in the brain indicates an attempt to counteract the toxic effects of exposure. Lastly, evidence in mammals has revealed increased selective constraints and limited baseline gene expression shifts in the brain relative to other tissues (Brawand et al., 2011; Fukushima & Pollock, 2020; Khaitovich et al., 2005). Such constraints could also influence the brain’s response to pesticide exposure, leading to less intense changes in gene expression relative to other tissues.

### Different tissues have varying capacities for pesticide detoxification

We detected the highest number of differentially expressed genes in Malpighian tubules. This organ likely accumulates higher levels of pesticides due to its primary excretory role analogous to the mammalian kidney and liver (Yang et al., 2022). Intriguingly, there was a general down-regulation of detoxification genes in Malpighian tubules. A comparable pattern was documented in the fruit fly, where fat body, another detoxification organ, showed a general down-regulation of detoxification genes after exposure to pesticides (Martelli et al., 2023). There are two possible explanations for these patterns. First, disturbances of the primary function of these tissues may compromise detoxification mechanisms after pesticide exposure over extended periods, potentially increasing susceptibility to additional stressors managed by the detoxification system (Siviter, Bailes, et al., 2021). Alternatively, the tissues may be reallocating resources to focus on the most pressing detoxification needs.

All three tissues exhibited changes in the expression of detoxification genes, although such changes were inconsistent. Clothianidin exposure disrupts detoxification activity without inducing uniform responses across the body. If clothianidin is partially metabolized before reaching specific tissues, the activation of different genes in distinct tissues could reflect stages of detoxification. However, previous reports suggest the detoxification system is always alert rather than activated by exposure (Bass et al., 2024). Accordingly, the three bumble bee cytochrome P450s most able to metabolize neonicotinoids constitute 33% of the total expression of the 51 cytochrome P450s in unexposed bumble bee brains (*X^2^ =* 21.5, *P* < 10^-6^). Therefore, changes in the expression of detoxification genes likely relate to processes affected in these tissues. For instance, glutathione S-transferases that were up-regulated in all three tissues participate in defense against oxidative stress in cells (Hong et al., 2024), which neonicotinoid pesticides can cause (Wang et al., 2018). The down-regulation of detoxification genes may also suggest resource redistribution, where certain detoxification processes are suppressed so the cells can allocate resources towards critical survival functions and maintaining essential metabolic activities under prolonged stress or damage (Lamichhane & Samir, 2023).

### Lowered energy metabolism may drive other metabolic changes

Disruption of specific cellular processes may result from the inhibition of energy metabolism, which we observed across all three tissues. Potentially, pesticide exposure causes a slowing down of the entire metabolism. Indeed, past studies have shown that pesticides cause mitochondrial depolarization (Moffat et al., 2015; Xu et al., 2022) and affect thermoregulation (Crall et al., 2018), navigation (Fischer et al., 2014), foraging activity (Muth & Leonard, 2019; Scholer & Krischik, 2014), and ovarian development, as shown here. Furthermore, alterations in the expression of genes linked to lipid metabolism and muscle functionality in the hind femur may influence energy storage, locomotive behavior, and colony development, all previously reported to be affected by neonicotinoids (Crall et al., 2018; Muth & Leonard, 2019; Siviter, Richman, et al., 2021). Beyond the direct effects of pesticides on tissues, some molecular changes from exposure may result from reduced metabolic rates and the redistribution of resources.

## Conclusions

Research on pesticide risks has traditionally focused on survival rates and select phenotypic traits. However, examining predetermined traits may capture a narrow range of pesticide exposure effects. Thus, pesticide risks could be either overlooked or overestimated, compromising the accuracy of safety assessment (EFSA, 2023). By contrast, tissue-specific transcriptomics allows simultaneously evaluating an exhaustive set of localized processes.

Our study highlights the complexity of pesticide impacts, demonstrating that different anatomical structures respond variably to exposure. Initial transcriptomic screenings can be a powerful tool for identifying which processes in which body parts are likely to be affected at the organismal level. This approach can identify specific traits for targeted assays, improving the detection of sub-lethal pesticide effects. Early studies in gene expression related to human diseases faced challenges in connecting changes in individual genes to tangible health outcomes (Hong et al., 2020). Similarly, while we cannot yet directly link gene expression changes to the ability of pollinators to reproduce or pollinate, transcriptomics can identify potential molecular biomarkers. For example, genes affected across various bumble bee tissues, such as the conserved pelota gene or those involved in the electron transport chain, may indicate pesticide impacts.

Integrating transcriptomics could greatly improve the precision of pesticide impact assessments, providing deeper insights into functional disruptions and sub-lethal effects caused by these chemicals and other human-induced stressors. Ultimately, such research can inform the development of more effective and eco-friendly pest management strategies, helping to conserve vital pollinator species and the ecosystems they sustain.

## Materials and Methods

### Microcolony setup and sampling

We used ten colonies of *Bombus terrestris audax* (Agralan Growers UK) to establish microcolonies, transferring the queen, existing brood and 20 workers to wooden boxes (30 cm wide, 20 cm long, 15 cm deep) separated into two equal-sized chambers (foraging and nesting area). Each colony received *ad libitum* 30% sucrose solution and organic honeybee-collected pollen (General Food Merchants LTD). We marked all 20 workers using water-based, non-toxic markers (POSCA, UK) and screened the colonies daily for the emergence of new workers. All queenright colonies were two weeks old when we started setting up the microcolonies. Six callow workers emerged within 24 hours and were used to assemble microcolonies. We obtained two microcolonies from each queenright colony, producing 20 microcolonies. We randomly assigned the control or clothianidin treatment to the two microcolonies from the same queenright colony. Each microcolony was kept in a single-chamber wooden box (12 cm wide, 12 cm long, 10 cm deep) and provided with organic pollen *ad libitum* and a single dose of nest substrate (5 parts organic pollen to one part 30% sucrose solution in a 2.5 cm Petri dish). Each microcolony received 10 mL of 30% control sucrose solution across two 5 ml syringes. We replaced the control syringes with two syringes containing pesticide solution after two days. We changed the pollen and sugar solution every 48 hours between 11:00 am and 1:00 pm. All colonies remained in darkness at 24 □, and we used red light during feeding and sampling. All sampling took place between 1:00 pm and 2:00 pm to minimize the effects of circadian rhythms on gene expression. We placed individual workers in 2 mL screw-cap cryovial tubes, rapidly submerged them in liquid nitrogen, and stored the samples at -80□.

### Pesticide solution preparation

We based the concentration of clothianidin on the residues found in pollen and nectar (Botías et al., 2015, 2015; David et al., 2016; Pohorecka et al., 2012; Rundlöf et al., 2015; Siviter et al., 2018). We prepared pesticide stock solution by dissolving clothianidin in acetone to a 0.5 mg/mL concentration and stored it in darkness at -20 □. Subsequently, we diluted the stock solutions using 30% sucrose solution to 5μg/L feeding solutions. Given that the weight of a liter of 30% sucrose solution is 1130 g, the final concentration was 4.4 ppb. To avoid pesticide degradation, we stored feeding solutions in darkness at 4 □.

### Tissue dissection, RNA extraction, library preparation and high throughput sequencing

Bumble bee abdomens were dissected in RNAlater. Ovaries of all bees were dissected and measured using millimeter paper. One worker per microcolony with the longest ovaries was recognized as dominant and excluded from sample processing for RNA-seq since significant ovarian development can lead to changes in bee physiology. We dissected Malpighian tubules in RNAlater, and brains and hind femurs on dry ice. We used three randomly selected non-dominant workers per microcolony and pooled candidate tissues for further steps. We homogenized the dissected tissues in 400 ml of TRIzol using FastPrep96 (45 seconds at 1800 RPM). We isolated RNA using chloroform and purified it with Genaxxon RNA Mini Spin Kit, applying DNase I on-column digestion. We checked RNA quality using TapeStation, and all samples had RIN scores > 8. We prepared cDNA libraries using the NEBNext® Ultra™ II Directional RNA Library Prep Kit for Illumina. We modified the standard protocol using one-third volumes of enzymes and buffers and 300 ng of total RNA input with 14 PCR cycles. The library size was quantified using TapeStation 2200 (Agilent, UK) and Qubit 2.0 fluorometer. Libraries were sequenced on NextSeq 500, generating a mean of 34.7 million 40 bp paired-end reads per sample (from 11 million to 45.8 million reads).

### Quality assessment of raw reads

We assessed the quality of raw reads using FastQC v0.11.9 (Andrews, 2019). To evaluate alignment qualities, we respectively aligned RNA-seq samples to the *B. terrestris* genome and transcriptome (Ensembl Metazoa release 52) using STAR v2.7 (Dobin et al., 2013) and kallisto v0.46 (Bray et al., 2016). We processed STAR alignments using the RNA-seq module of Qualimap v2.2.1 (Okonechnikov et al., 2016). Out of the 120 samples, two failed during sequencing and another two failed the quality-control checks due to poor read alignment (< 60% reads aligned by kallisto, Conesa et al., 2016). We, therefore, removed two samples from each group (matchins samples from failed colonies or removing samples with the lowest number of reads), retaining eight replicates for all combinations of tissues and treatments to ensure balanced statistical testing.

### Differential gene expression analysis

We separately summarized kallisto pseudo-aligned transcript abundances to the gene level with tximport v1.14 (Soneson et al., 2016) for each tissue and pesticide using transcript-to-gene tables retrieved with biomaRt v2.48.3. We filtered out genes with relatively low expression, choosing a cut-off of 20 reads in at least 50% of samples and at least 80 reads. We applied treatment-aware filtering as implemented in *filterByExpr* function from EdgeR v3.32.1. Each tissue/pesticide combination was analyzed separately against the corresponding control because of large differences among biological replicates between tissues. To detect differentially expressed genes between exposure treatments and the control, we applied Wald tests on median-of-ratios normalized counts in DESeq2 v1.32.0 (Love et al., 2014). We included treatment and source colony as factors in the model formula. We report genes as differentially expressed using a significance cut-off of 0.05 after false discovery rate adjustment of the Wald test *P*-values (FDR).

### Weighted Gene Co-expression Network Analysis (WGCNA)

We constructed a multi-tissue co-expression (consensus) network to decipher modules of genes conserved between tissues and to ask whether some of these modules are affected by clothianidin. This network was built on genes that were active in all three tissues (n = 7069) using all samples (n = 48). Performing this analysis allowed us to detect alterations in gene expression that are shared across tissues after exposure to clothianidin. We first created a shared DESeq2 object for all samples (n = 48) and applied median-of-ratio normalization in DESeq2 to account for potential variations in the library composition. We then applied the variance stabilizing transformation (VST) to ensure that the variance was not mean-dependent and constructed a weighted gene co-expression network using WGCNA v.1.72 R package. We applied a soft-thresholding power of 25 to the correlation matrix based on the scale-free topology fit index of 0.8. Each correlation value was raised to the power of 25, effectively emphasizing the stronger correlations between genes. Through this approach, we prioritized and focused on the more substantial, biologically meaningful correlations. After constructing the network, we tested for correlation with exposure treatment: exposed vs. unexposed using eigengenes and applying Pearson correlation and Benjamini-Hochberg *P*-value adjustment for multiple testing. A module eigengene is the first principal component of a given module and can be considered a representative expression profile for that module. We considered modules as significantly associated with the exposure treatment by applying a cut-off of 0.05 to Pearson correlation adjusted *P*-values. To visualize the network, we focused on the top 5 most robust connections and adopted an upper quantile of 0.25 for the strength of correlation, ensuring clarity and emphasis on the most relevant connections within the network.

### Unsupervised clustering of gene expression patterns across tissues

We used unsupervised clustering to detect patterns of co-expression of genes across tissues of *B. terrestris*. Performing this analysis allowed us to detect relative differences in baseline gene expression between tissues. We used exclusively control samples (n = 24). We applied the median-of-ratios normalization implemented in DESeq2 and used NbClust v.3.0.1 R library to perform k-means clustering on all samples together. Based on the Calinski-Harabasz index, we established six gene co-expression clusters for all tissues with higher relative expression in each tissue and each pair of tissues.

### Gene Ontology and KEGG enrichment analysis

To test which Gene Ontology (GO) terms were overrepresented in response to the treatments, we performed gene ontology enrichment analysis using *g:GOSt* option of gprofiler2 v0.2.1 (Kolberg et al. 2020). We sorted differentially expressed genes for each treatment by *P*-values and set the custom background genes to all genes expressed in control samples for each tissue separately. We report GO terms as enriched by applying the software’s internal correction for multiple testing g:SCS threshold of 0.05 derived from Fisher’s exact test.

### Detoxification genes detection

To identify genes likely involved in detoxifying pesticides, we searched the proteome for the following Pfam domains: PF00067 (cytochrome P450), PF00664, PF00005 (ABC-transporters), PF02798, PF00043 (glutathione S-transferases), PF00201 (UDP-glucoronosyl and UDP-glucosyl transferases), and PF00135 (carboxylesterases) and extracted the recognized amino acid sequences using HMMER v3.1b2 applying the --cut_ga option, which removes domains with conditional E-values greater than the internally established threshold. We further manually curated the lists of candidate genes using BLAST.

### Modelling ovary length after exposure to clothianidin

We chose ovary length as the phenotypic trait to illustrate if molecular-level metabolic alterations caused by chronic clothianidin exposure have consequential effects on observable traits. We obtained measurements for 49 bees exposed to clothianidin and 55 for the control. The uneven number of individuals measured was due to higher mortality in clothianidin-exposed microcolonies. Given that the data exhibited a zero-inflated distribution along with an otherwise normal distribution, a two-part mixed-effects model designed for semi-continuous data was employed using the GLMMadaptive v. 0.9-1 package in R with a custom distribution family function to account for the binary and normally-distributed fractions of the data. The treatment was incorporated as a fixed effect, while the source colony was treated as a random effect. The random effect was applied to both the binomial and normally-distributed components of the model. To validate the model’s accuracy, we generated simulated data over 25 iterations and performed a posterior predictive check. This procedure allowed us to assess how well the fitted model could reproduce the characteristics of the observed data.

## Data and code availability

Sequence data underlying this work are in the National Center for Biotechnology Information Short Read Archive, accessions PRJNA1076820 & (available upon publication). Analysis scripts will be available on GitHub at https://github.com/wurmlab/wurmlab/2023-Bumblebee-tissue-specific-effects-of-clothianidin-insecticide. During review, analysis scripts can be found at https://www.dropbox.com/scl/fo/tzcm5i8iuwmdy1v53fo7p/AKyJDnTHkaI6kCmBHorfI?rlkey=8762kl8wyg4gwulwqllrl71uz&st=pcun5pri&dl=0

## Acknowledgments

We were funded by the Natural Environment Research Council (NE/L00626X/1 and NE/S007229/1), the European Commission (H2020-MSCA-IF-2018-840185), and the Biotechnology and Biological Sciences Research Council (BB/T015683/1). This research used the Apocrita HPC facility, supported by QMUL Research-IT (http://doi.org/10.5281/zenodo.438045).

## Supplementary Materials

**Supplementary Figure 1.** Changes in ovarian growth after chronic clothianidin exposure. **A.** Ovariole length [mm] in bumble bee workers exposed to the control and clothianidin. **B.** Histograms of the ovariole lengths in bumble bee workers; many of the clothianidin-exposed bees did not develop ovaries. C. Cumulative density function (CDF) of the ovariole length simulated with the fit model over 15 iterations. The fit data follows the simulated trend showing an appropriate model fit.

**Supplementary Figure 2.** Co-expression (k-means) clusters of genes. The lines show the relative trend of mean scaled expression of genes classified into each of the clusters in different tissues.

**Supplementary Table 1.** Detoxification genes identified in *B. terrestris* separated into families and subfamilies. Genes were assigned to one of the co-expression (k-means) clusters. Arrows indicate trends in differential expression, colors indicate tissues.

