## Supplementary Table 1 for "A neonicotinoid pesticide causes tissue-specific gene expression changes in bumble bees"

**Supplementary Table 1.** Detoxification genes identified in *B. terrestris* separated into families and subfamilies. Genes were assigned to one of the co-expression (k-means) clusters. Arrows indicate trends in differential expression, colors indicate tissues.

| Gene ID | Family | Subfamily | k-mean Cluster | Average expression in control samles: Malpighian tubules | Brain | Hind femur | SD | Differential expression |
|---|---|---|---|---|---|---|---|---|
| LOC100650427 | Cytochromes P450 | CYP9Q | NA | 1.07 | 1.58 | 0.49 | 0.55 | |
| LOC100648995 | | | I | 85.72 | 6.02 | 7.62 | 45.56 | |
| LOC100643678 | | | III | 1.82 | 32.10 | 1,155.59 | 657.56 | |
| LOC105666367 | | | III | 1.72 | 25.18 | 1,011.85 | 576.55 | ↓ (Hind femur) |
| LOC105666366 | | | I | 130.40 | 2.93 | 7.61 | 72.28 | |
| LOC100642936 | | | III | 6,626.26 | 653.84 | 17,689.21 | 8643.52 | ↓ (Malpighian tubules) ↑ (Hind femur) |
| LOC105666343 | | | I | 357,999.72 | 165.14 | 10,886.91 | 203571.38 | ↓ (Malpighian tubules) |
| LOC100647803 | | | I | 2,963.86 | 4.55 | 135.55 | 1672.03 | ↓ (Malpighian tubules) |
| LOC100642816 | | | I | 1,700.69 | 2.83 | 60.69 | 963.99 | |
| LOC100649110 | | | I | 418.31 | 3.83 | 51.35 | 226.83 | ↓ (Malpighian tubules) |
| LOC100648391 | | | IV | 580.42 | 7,121.61 | 4,233.88 | 3278.06 | ↑ (Brain) |
| LOC100651291 | | | II | 25.88 | 53.00 | 68.02 | 21.36 | |
| LOC100647234 | | | I | 7,045.06 | 431.53 | 2,386.48 | 3397.62 | ↑ (Malpighian tubules) |
| LOC100647041 | | | I | 31.43 | 47.20 | 46.91 | 9.02 | ↓ (Brain) |
| LOC100646434 | | | I | 25,985.22 | 2,015.00 | 3,331.41 | 13475.28 | |
| LOC110119312 | | | NA | 1.52 | 0.45 | 0.78 | 0.55 | |
| LOC100648257 | | | II | 4,033.99 | 36.05 | 3,016.63 | 2077.76 | ↑ (Malpighian tubules) |
| LOC100648029 | | | I | 25,175.52 | 178.22 | 6,982.14 | 12923.86 | ↓ (Malpighian tubules) |
| LOC100649469 | | | I | 192.31 | 2.53 | 113.17 | 95.32 | ↓ (Hind femur) |
| LOC100646677 | | | I | 30,072.40 | 892.01 | 4,017.09 | 16021.55 | ↑ (Hind femur) |
| LOC100647549 | | | II | 6,819.88 | 1,494.93 | 2,345.15 | 2860.69 | ↑ (Hind femur) |
| LOC100647785 | | | IV | 7,819.92 | 20,880.90 | 7,232.50 | 7715.93 | |
| LOC100648939 | | | NA | 0.12 | 0.08 | 0.00 | 0.06 | |
| LOC100651693 | | | I | 3,888.92 | 437.11 | 385.09 | 2008.09 | ↓ (Malpighian tubules) |
| LOC100647886 | | | NA | 4.52 | 1.86 | 2.41 | 1.40 | |
| LOC100647566 | | | IV | 24,374.79 | 28,263.32 | 16,302.36 | 6101.22 | ↑ (Brain) |
| LOC100648545 | | | I | 1,207.48 | 163.29 | 720.66 | 522.49 | ↑ (Malpighian tubules) |
| LOC100649760 | | CYP336A | I | 33,137.53 | 4,231.76 | 1,160.21 | 17642.41 | |
| LOC100649871 | | | VI | 11,344.01 | 11,523.13 | 878.31 | 6094.74 | |
| LOC100649988 | | | IV | 2,311.69 | 10,411.61 | 1,048.58 | 5080.53 | ↑ (Hind femur) |
| LOC100649441 | | | I | 401.61 | 100.03 | 16.93 | 202.42 | |
| LOC100647251 | | CYP9Q | I | 74,113.93 | 25,372.15 | 5,886.28 | 35143.69 | |
| LOC100647368 | | | IV | 2,919.37 | 19,215.65 | 2,939.59 | 9402.83 | |
| LOC100650309 | | | VI | 117.92 | 144.49 | 55.31 | 45.79 | |
| LOC100652170 | | Clan 4 | III | 6,653.49 | 297.61 | 11,322.38 | 5533.85 | ↓ (Malpighian tubules) |
| LOC100647609 | | | I | 193.47 | 4.58 | 4.36 | 109.12 | |
| LOC100644743 | | | III | 313.99 | 449.42 | 713.42 | 203.13 | ↑ (Brain) |
| LOC100648944 | | | III | 173.19 | 224.73 | 356.71 | 94.65 | |
| LOC100651255 | | | I | 6.74 | 5.06 | 3.96 | 1.40 | |
| LOC100647578 | | Clan 2 | III | 32.57 | 1,209.70 | 9,044.64 | 4898.80 | |
| LOC100648323 | | | II | 90.10 | 73.20 | 82.86 | 8.48 | |
| LOC100651206 | | | I | 49.65 | 67.18 | 29.53 | 18.84 | |
| LOC100650054 | | | IV | 7.61 | 74.59 | 43.00 | 33.51 | |
| LOC100642897 | | | III | 46.94 | 142.28 | 892.13 | 462.91 | |
| LOC100642619 | | | IV | 53.50 | 52.22 | 49.19 | 2.21 | |
| LOC100645445 | | Mitochondrial | III | 12.69 | 14.57 | 162.40 | 85.90 | |
| LOC100646342 | | | III | 1.16 | 2.94 | 13.90 | 6.90 | |
| LOC100645879 | | | NA | 2.13 | 3.60 | 0.30 | 1.65 | |
| LOC100649492 | | | IV | 125.30 | 302.84 | 136.66 | 99.39 | |
| LOC100649449 | | | I | 6,033.02 | 84.43 | 323.79 | 3367.45 | ↑ (Malpighian tubules) |
| LOC100644009 | | | III | 140.90 | 116.81 | 514.14 | 222.77 | |

| Gene ID | Family | Subfamily | k-mean Cluster | Average expression in control samles: Malpighian tubules | Brain | Hind femur | SD | Differential expression |
|---|---|---|---|---|---|---|---|---|
| LOC100645338 | ABC-transporters | ABCC | III | 78.74 | 296.12 | 4,321.71 | 2,389.40 | |
| LOC100651734 | | | V | 316.77 | 1,574.08 | 1,411.17 | 683.75 | |
| LOC100645209 | | | I | 53,654.88 | 3,753.00 | 3,355.23 | 28,926.37 | ↓ (Malpighian tubules) ↑ (Brain) |
| LOC100650642 | | | I | 32,402.63 | 965.43 | 1,317.43 | 18,049.52 | ↑ (Malpighian tubules) ↑ (Hind femur) |
| LOC100644974 | | | I | 6,034.75 | 1,995.62 | 971.36 | 2,677.11 | |
| LOC100646583 | | | I | 10,644.60 | 3,710.57 | 1,547.96 | 4,752.31 | |
| LOC100651558 | | | IV | 481.27 | 655.13 | 506.73 | 93.90 | |
| LOC100644751 | | | I | 775.59 | 75.99 | 90.17 | 399.88 | |
| LOC105666350 | | | I | 4,508.59 | 1,109.20 | 565.65 | 2,136.90 | |
| LOC100649886 | | ABCB | III | 186.67 | 275.66 | 418.57 | 116.99 | ↓ (Brain) |
| LOC100650335 | | | II | 720.28 | 334.39 | 675.96 | 211.17 | |
| LOC100652256 | | | I | 3,242.28 | 1,510.81 | 1,505.63 | 1,001.16 | |
| LOC100649698 | | | VI | 1,615.16 | 1,188.62 | 780.60 | 417.31 | |
| LOC100650108 | | | III | 9,775.62 | 8,557.92 | 42,205.92 | 19,084.88 | ↓ (Malpighian tubules) ↑ (Brain) |
| LOC100646327 | | ABCD | III | 1,045.92 | 976.32 | 1,181.25 | 104.21 | |
| LOC100643487 | | | III | 1,080.79 | 670.81 | 2,327.40 | 862.79 | ↑ (Brain) |
| LOC100650667 | | ABCF | III | 1,071.08 | 749.59 | 1,173.13 | 221.04 | |
| LOC100651511 | | | II | 1,442.92 | 1,053.01 | 1,679.55 | 316.38 | |
| LOC100648932 | | | II | 805.05 | 476.53 | 843.96 | 201.84 | ↓ (Malpighian tubules) |
| LOC100644595 | | ABCE | II | 1,160.93 | 655.43 | 1,014.33 | 260.07 | |
| LOC100644723 | | ABCG | II | 1,203.04 | 175.99 | 1,632.77 | 748.52 | |
| LOC100644848 | | | IV | 952.75 | 2,657.58 | 1,488.50 | 871.80 | ↑ (Brain) |
| LOC100644985 | | | III | 14.95 | 63.41 | 989.83 | 549.39 | ↑ (Hind femur) |
| LOC100644403 | | | III | 87.15 | 313.88 | 2,539.04 | 1,354.90 | |
| LOC100644744 | | | III | 2,542.74 | 1,703.98 | 6,325.25 | 2,461.94 | |
| LOC100644506 | | | V | 317.84 | 416.60 | 315.35 | 57.75 | ↑ (Hind femur) |
| LOC100646359 | | | III | 493.48 | 203.73 | 2,018.46 | 974.92 | |
| LOC100644523 | | | I | 1,068.79 | 413.73 | 465.17 | 364.26 | |
| LOC100648854 | | | VI | 318.47 | 185.21 | 167.64 | 82.48 | |
| LOC100644150 | | | I | 1,347.04 | 34.66 | 44.79 | 754.80 | |
| LOC100644270 | | | I | 4,616.32 | 4.15 | 108.64 | 2,633.19 | |
| LOC100646233 | | | I | 11,070.93 | 349.74 | 384.41 | 6,179.90 | ↑ (Malpighian tubules) |
| LOC100643668 | | | III | 18.45 | 21.50 | 105.81 | 49.58 | |
| LOC100648840 | | | III | 14.15 | 13.67 | 27.12 | 7.63 | |
| LOC100645470 | | | III | 583.84 | 871.69 | 6,043.42 | 3,072.37 | ↓ (Malpighian tubules) |
| LOC100649393 | | | III | 346.07 | 2,077.54 | 8,574.14 | 4,337.91 | |
| LOC100645221 | | | III | 51.94 | 1,307.66 | 607.40 | 629.25 | ↑ (Hind femur) |
| LOC100650392 | | ABCA | III | 576.12 | 1,807.52 | 4,523.07 | 2,019.45 | |
| LOC100645517 | | | VI | 3,320.10 | 2,404.76 | 1,779.50 | 774.84 | |
| LOC100647302 | | | III | 1,475.95 | 1,558.80 | 1,461.28 | 52.58 | |

| Gene ID | Family | Subfamily | k-mean Cluster | Average expression in control samles: Malpighian tubules | Brain | Hind femur | SD | Differential expression |
|---|---|---|---|---|---|---|---|---|
| LOC105667063 | Glutathione S-transferases | | II | 26.04 | 20.74 | 35.59 | 7.53 | |
| LOC105667074 | | | VI | 234.91 | 345.22 | 160.46 | 92.96 | ↑ (Malpighian tubules) ↑ (Brain) |
| LOC105667024 | | | II | 266.06 | 206.70 | 244.08 | 30.01 | ↑ (Brain) |
| LOC100644200 | | | VI | 139.64 | 105.86 | 61.89 | 38.99 | |
| LOC100645775 | | | III | 12.29 | 49.68 | 414.22 | 222.05 | ↓ (Brain) ↑ (Hind femur) |
| LOC100646009 | | | III | 7,444.23 | 4,102.78 | 13,198.41 | 4600.84 | ↓ (Malpighian tubules) |
| LOC100646355 | | | III | 9.49 | 132.89 | 621.10 | 323.43 | |
| LOC100651606 | | | I | 489.05 | 180.87 | 251.65 | 161.42 | |
| LOC100650722 | | | I | 58,410.62 | 25,036.96 | 20,278.55 | 20778.59 | ↓ (Malpighian tubules) ↑ (Hind femur) |
| LOC100651245 | | | IV | 296.83 | 1,323.69 | 61.03 | 671.36 | |
| LOC100650878 | | | III | 3.98 | 15.55 | 54.75 | 26.61 | |
| LOC100651035 | | | VI | 3,378.75 | 1,927.94 | 1,993.09 | 819.47 | ↓ (Malpighian tubules) |
| LOC100647524 | UDP-glycosyltransferases | UDP-glucuronosyltransferases | I | 34,901.55 | 96.39 | 902.50 | 19866.15 | |
| LOC100647883 | | | VI | 36,619.51 | 18,815.78 | 2,091.08 | 17267.02 | ↓ (Malpighian tubules) ↑ (Brain) |
| LOC100646290 | | | III | 0.11 | 1.80 | 48.00 | 27.17 | |
| LOC100649407 | | | V | 3,345.99 | 3,079.53 | 5,730.54 | 1459.73 | |
| LOC100651802 | | | IV | 299.74 | 553.39 | 479.96 | 130.52 | |
| LOC100642358 | | | II | 3.09 | 3.27 | 33.90 | 17.74 | |
| LOC100647646 | | | I | 739.40 | 38.36 | 45.82 | 402.61 | ↓ (Malpighian tubules) |
| LOC100648003 | | | VI | 37.13 | 22.60 | 2.15 | 17.57 | |
| LOC100645384 | | UDP-glucosyltransferases | I | 13,262.31 | 6,218.22 | 2,719.48 | 5369.85 | |
| LOC100645683 | | | IV | 997.60 | 7,195.05 | 3,425.10 | 3122.86 | ↓ (Malpighian tubules) |
| LOC100645567 | | | III | 106.77 | 383.79 | 7,219.32 | 4028.85 | |
| LOC100645199 | Carboxylesterases | | III | 368.20 | 727.83 | 724.88 | 206.79 | ↑ (Malpighian tubules) ↑ (Hind femur) |
| LOC100652311 | | | VI | 10.76 | 2.52 | 1.49 | 5.08 | |
| LOC100652071 | | | III | 1.42 | 1.55 | 1.03 | 0.27 | |
| LOC100647870 | | | III | 799.37 | 648.87 | 4,732.55 | 2315.49 | ↓ (Brain) |
| LOC100652190 | | | I | 16.44 | 1.02 | 0.29 | 9.12 | |
| LOC100652269 | | | I | 171.66 | 47.77 | 16.52 | 82.05 | ↑ (Malpighian tubules) |

Malpighian tubules
Hind femur
Brain
