## Supplementary figures and images for "A neonicotinoid pesticide causes tissue-specific gene expression changes in bumble bees"

### Supplementary Figure 1

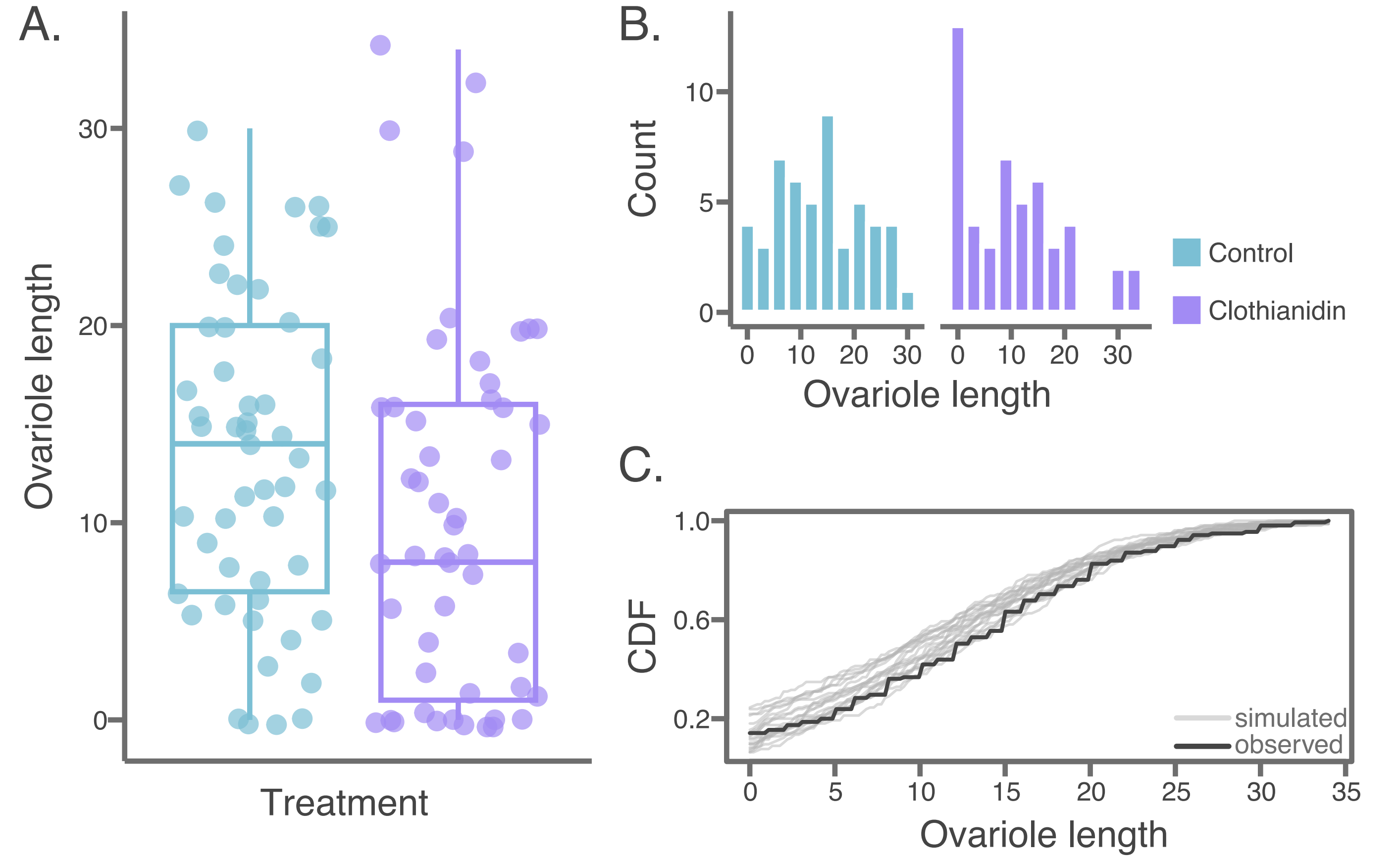

### Supplementary Figure 2

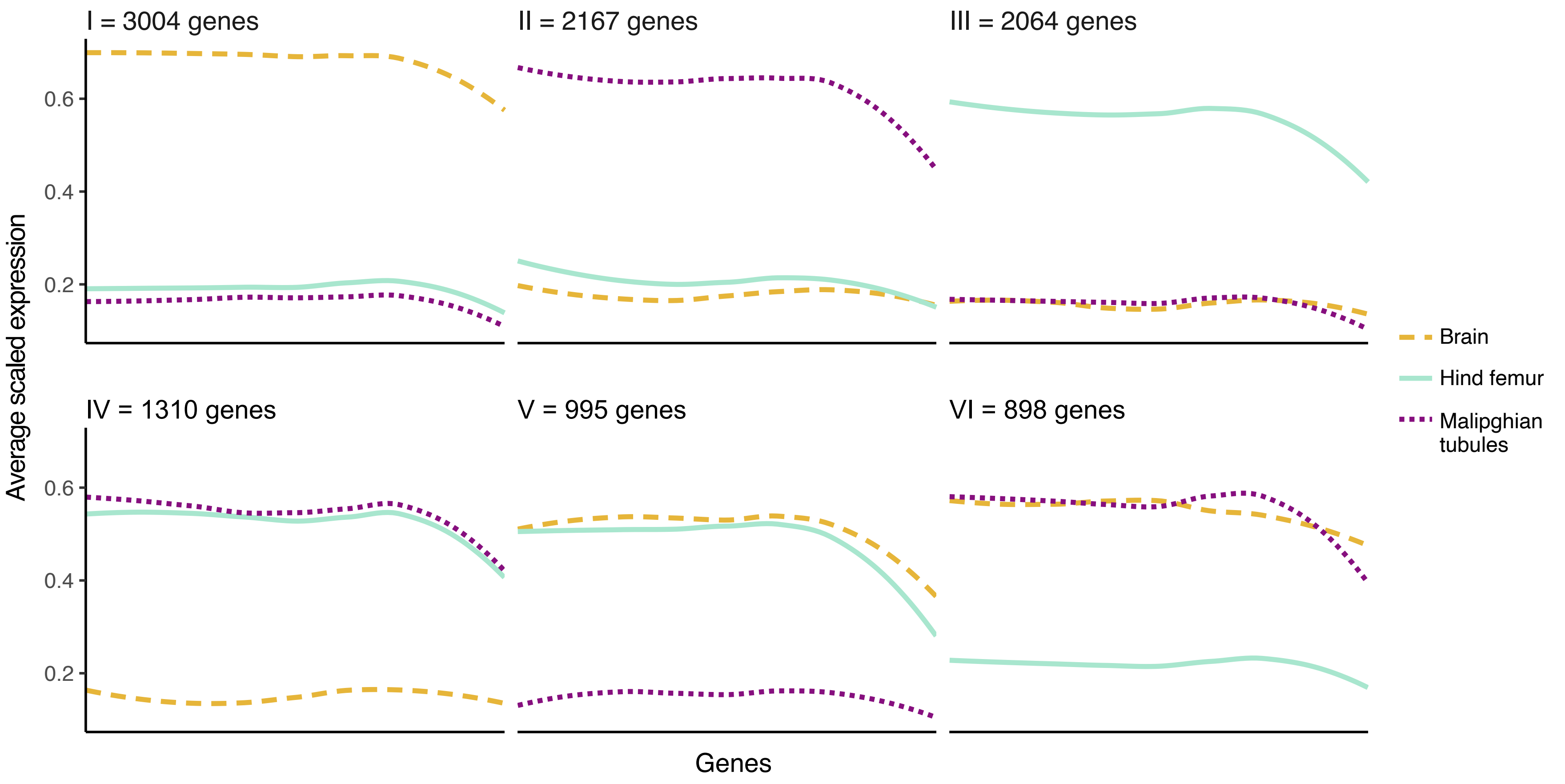
